# Selective autophagy, lipophagy and mitophagy, in the Harderian gland along the estrous cycle: a potential retrieval effect of melatonin

**DOI:** 10.1101/652222

**Authors:** Marina García-Macia, Adrián Santos-Ledo, Beatriz Caballero, Adrián Rubio-González, Beatriz de Luxán-Delgado, Yaiza Potes, Susana Rodríguez-González, José Antonio Boga, Ana Coto-Montes

**Affiliations:** Institute of Cellular Medicine, Newcastle University, NE2 4HH, Newcastle Upon Tyne, UK; Institute of Genetic Medicine, Newcastle University, NE2 4HH, Newcastle Upon Tyne, UK; Departamento de Morfología y Biología Celular, Área de Biología Celular, Facultad de Medicina, Universidad de Oviedo, Julián Clavería s/n, 33006, Oviedo, Spain; Servicio de Microbiología, Hospital Universitario Central de Asturias, Celestino Villamil s/n, 33006 Oviedo, Spain

**Keywords:** autophagy, lipophagy, mitochondria, oxidative stress, sexual hormones, Harderian gland

## Abstract

Sexual dimorphism has been reported in many processes. However, sexual bias in favor of the use of males is very present in science. One of the main reasons is that the impact of hormones in diverse pathways and processes such as autophagy have not been properly addressed *in vivo.* The Harderian gland is a perfect model to study autophagic modulation as it exhibits important changes during the estrous cycle. The aim of this study is to identify the main processes behind Harderian gland differences under estrous cycle and their modulator. In the present study we show that redox-sensitive transcription factors have an essential role: NF-κB may activate SQSTM1/p62 in estrus, promoting selective types of autophagy: mitophagy and lipophagy. Nrf2 activation in diestrus, leads the retrieval phase and restoration of mitochondrial homeostasis. Melatonin’s receptors show higher expression in diestrus, leading to decreases in pro-inflammatory mediators and enhanced Nrf2 expression. Consequently, autophagy is blocked, and porphyrin release is reduced. All these results point to melatonin as one of the main modulators of the changes in autophagy during the estrous cycle.

## INTRODUCTION

Sex bias is still a clear issue in biomedical research, although sexual dimorphism has been described in many cellular processes under pathological and physiological conditions. Sex-dependent differences in the activation of autophagy have been reported mainly *in vitro* (Du et al. 2009; Vega-Naredo et al. 2009). It has been reported that hormonal changes stimulate autophagy in mammary gland, but just using *in vitro* models (Zielniok et al. 2014; Zielniok et al. 2017). Androgens also modulate autophagy in flank organ, which is a sebaceous gland, *in vivo* (Coto-Montes et al. 2009). However, how autophagy is modulated under sexual hormone variations, as during the estrous cycle, is poorly understood.

The Syrian hamster Harderian gland (HG) is a tubule-alveolar orbital gland formed by two main types of cells (type I and II) that secretes lipids favoring lubrication of the cornea (Sakai 1981). The HG has other functions including the production of pheromones (Payne et al. 1979), functions related to the pineal-gonadal axis (Hoffman et al. 1985), and synthesis of indolamines (Menendez-Pelaez et al. 1993). HG exhibits marked sexual differences in relation to cell type and porphyrin production. In previous studies, we have demonstrated alterations of female gland activity during the estrous cycle (Garcia-Macia et al. 2014). Interestingly, the estrus phase presents the highest porphyrin–production activity, which has been associated with a prominent oxidative damage to proteins, together with a lower total antioxidant activity and higher autophagy, compared to the diestrus phase (Garcia-Macia et al. 2014; Vega-Naredo et al. 2009). By contrast, the diestrus phase shows a lower porphyrins production causing reduced oxidative stress levels, a decrease in proteolytic activities and a blockage of autophagy due to activation of mTOR (Garcia-Macia et al. 2014). Morphological changes also occur during the estrous cycle, the Type II cells of the Harderian gland are more abundant during estrus.

The main difference between Type I and II cells is the size of the lipid droplets (LD). Type II cells present bigger and more abundant LDs, suggesting that the LDs are active organelles in the HG. Previously, it has been reported that autophagy controls the mobilization of LDs to lysosomes (Singh and Cuervo 2012; Singh et al. 2009). This new type of LD degradation is called lipophagy (Singh and Cuervo 2012; Singh et al. 2009). Although it was first discovered in liver, it has also been observed in several cell types, where it acts as an energy source with the peculiar capability to provide large amounts of free fatty acids in a short time (Singh and Cuervo 2012; Singh et al. 2009). Turnover of endogenous lipid stores is essentially developed by Lysosomal acid lipase (LAL) (Duta-Mare et al. 2018; Pearson et al. 2014), which could be considered an essential lipophagy marker. However, the presence of lipophagy in HG is still unconfirmed, even though lipid droplets are indispensable components of glandular cells.

The pineal melatonin is the endogenous synchronizer of the circadian rhythms in organisms, mainly of those related to the control of seasonal reproductive phenomena (Reiter 1980). Likewise, melatonin and its metabolites are well-known antioxidants (Negi et al. 2011). Moreover, melatonin is able to maintain mitochondrial homeostasis (Coto-Montes et al. 2012; Tamarindo et al. 2019), as it increases the activity of the mitochondrial respiratory complexes I and IV, which result in increased ATP production (Martin et al. 2000). Melatonin plays and important role modulating the morphology and physiology of the HG (Menendez-Pelaez et al. 1993; Vega-Naredo et al. 2012). Melatonin secretion depends on the phase of the estrous cycle (Ozaki et al. 1978) and it is diminished by gonadal steroid hormones (Rato et al. 1999).

Based on these findings and considering the observed oscillations in autophagy activity as well as Type II cell abundance along the estrous cycle (Garcia-Macia et al. 2014), the aim of this study was to identify the main modulators behind autophagic changes in the estrous cycle and the involvement of the lipophagy.

## MATERIALS AND METHODS

### Animals

Eight-week-old female Syrian hamsters *(Mesocricetus auratus)* (Harlan Interfauna Ibérica, Barcelona, Spain) were housed 2 per cage under long days with a 14:10 light:dark cycle (lights on daily from 07:00 to 21:00) at 22 ± 2°C (n=8 per experimental condition). Hamsters received water and a standard pellet diet *ad libitum*. The Oviedo University Local Animal Care and Use Committee approved the experimental protocols (reference 33443591). All experiments were carried out according to the Spanish Government Guide and the European Community Guide for Animal Care (Council Directive 86/609/EEC).

All animals were monitored daily by vaginal smears to determine their reproductive phase (proestrus, estrus, metestrus, diestrus) according to the method of Orsini (Orsini 1961), during 3 consecutive cycles. For studying opposite phases, we collected females during the estrus and diestrus phases because these phases present the highest plasmatic differences in estrogen concentration during the estrous cycle (Lisk 1985). After determination of the specific estrous phase, animals were sacrificed, and the hamster Harderian glands (HGs) were immediately removed, frozen in liquid nitrogen, and stored at −80°C until performing of the experiments, HGs were used directly for lipid droplet isolation.

### Isolation of proteins

HGs (0.1 g) were homogenized using a Polytron homogenizer at 4°C in 1 ml of lysis buffer (50 mM Tris/HCl, 150 mM NaCl at pH 7.4 and protease and phosphatase inhibitors). The tissue homogenates were then centrifuged for 6 min at 3000 rpm at 4°C. The supernatants were collected and centrifuged again under the same conditions. The protein concentration of the supernatants was measured by the method of Bradford (Bradford 1976).

### Isolation of lipid droplet (LD) fractions

LD fraction from Harderian gland was isolated following the protocol for brown adipose tissue with few alterations (Martinez-Lopez et al. 2016). Tissues were homogenized in 0.25 M sucrose and centrifuged at 6,800 g for 5min/4°C. Supernatants including the fatty layer were centrifuged at 17,000 g/10min/4°C to eliminate unwanted cellular fractions. Supernatant from the 17,000 g spin was adjusted to 20% sucrose and centrifuged in a discontinuous sucrose density gradient at 27,000 g for 30 min at 4°C. LD fractions were delipidated using successive washes in acetone and ether and solubilized in 2% SDS for immunoblotting.

### Immunoblotting

Protein samples were prepared in western-blotting sample buffer (65.8 mM Tris-HCl, pH 6.8, 2.1% SDS, 26.3% (w/v) glycerol, 0.01% Bromophenol Blue). SDS-polyacrylamide gels were run and analyzed as previously described (Garcia-Macia et al. 2011; Vega-Naredo et al. 2012). Primary antibodies applied were as follows: nuclear factor erythroid 2-related factor 2 (Nrf2), superoxide dismutase 2 (SOD2), retinoid-related orphan receptor alpha (RORα) and melatonin receptor 1A (MT1) from Santa Cruz Biotechnology (Santa Cruz, CA, USA), adipose triglyceride lipase (ATGL), beclin 1, hormone-sensitive lipase (HSL), microtubule-associated protein 1A/1B-light chain 3 (LC3), nuclear factor kappa-B p65 subunit (NF-κB p65), nuclear factor kappa-B p65 subunit phosphorylated at Serine 536 (Phospho-NF-κB p65 [Ser536]) and sequestosome-1 (SQSTM1/p62) from Cell Signaling Technology (Boston, MA, USA), glyceraldehyde 3-phosphate dehydrogenase (GAPDH) and lysosomal acid lipase (LAL) antibody from Abcam (Cambridge, UK), lysosomal-associated membrane protein 1 (Lamp1) from Developmental Studies Hybridoma Bank (Iowa City, Iowa, USA); Bcl2/Adenovirus E1B 19 kDa and protein-interacting protein 3-like (BNIP3L/NIX) from Sigma-Aldrich (St. Louis, MO, USA), perilipin (PLIN) 2 (Progen Biotechnik, Heidelberg, DE) and PLIN3 (ProSci Inc, Poway, CA, US). Primary antibodies were mostly diluted 1:1000 in blocking buffer, except for Phospho-NF-κB p65 (Ser536), which was diluted 1:500, and Nrf2, which was diluted 1:250. Goat β-actin antibody (Santa Cruz Biotechnology, Inc.) diluted at 1:1000 was always assayed as a loading reference. All the primary antibodies have been previously validated (Garcia-Macia et al. 2014; Klionsky et al. 2016; Vega-Naredo et al. 2012; Vega-Naredo et al. 2009). After washing in TBS-T (20 mM Tris-HCl, 150 mM NaCl, pH 7.4 and 0.05% Tween-20), the membranes were then incubated with the corresponding horseradish peroxidase-conjugated secondary antibody: anti-rabbit for most primaries, but anti-mouse for β-actin, or antirat for Lamp1 or anti-guinea pig for the PLIN2 (Santa Cruz Biotechnology, Inc.) diluted 1:2500. Binding of antibodies to their antigens was detected using the Western Blotting Luminol Reagent (sc-2048; Santa Cruz Biotechnology, Inc.) according to the manufacturer’s protocol. Negative controls were performed with either no primary or no secondary antibodies. No bands were detected in any case. The results were calculated from at least three separate experiments for each antibody and were normalized to actin. Band intensity was quantified using the Quantity One 1D analysis software v. 5.5.1. (Bio-Rad Laboratories Inc., Hercules, CA, USA).

### Fluorescence microscopy

HGs were fixed overnight in 4% paraformaldehyde (PFA) solution at 4°C. After three washes in PBS, HGs were cryoprotected first overnight at 4°C in 15% sucrose and then 4 hours at room temperature in 30% sucrose. HGs were oriented and embedded in OCT and then frozen in liquid nitrogen. 10 μm sections were obtained in a cryostat. Sections were rinse in PBS and then blocked with 3% horse serum, 1% BSA in 1x PBS containing 0.4% triton X-100 for 1 hour at room temperature, then they were incubated with primary were incubated with primary antibodies Lamp1 (Developmental Studies Hybridoma Bank, Iowa City, Iowa, USA) and LC3 (Cell Signalling Technology, Boston, MA, USA) and secondary antibodies anti-rat and anti-rabbit, respectively (Alexa Fluor 647 conjugated, Invitrogen (Carlsbad, Ca, USA)). For lipid droplet (LD) detection, sections were incubated with BODIPY 493/503 for 20min at RT (Molecular Probes Inc., Eugene, OR, USA). Mounting medium contained DAPI (4’, 6-diamidino-2-phenylindole) to visualize the nucleus (Molecular Probes Inc., Eugene, OR, USA). Negative controls were performed with either no primary or no secondary antibodies. No staining was detected in any case. Images were acquired on a Nikon A1R confocal inverted (Nikon Instruments INC., NY, USA) using X100 objective/1.4 numerical aperture. Images were acquired at similar exposure times in the same imaging session. Image slices of 0.2μm thickness were acquired and deconvolved using the Huygens (Huygens Essential, Hilversum, The Netherlands) acquisition/analysis software. Quantification was performed in deconvolved images after appropriate thresholding using the ImageJ software (NIH) (Schneider et al. 2012) in a minimum of 30 acini from at least 3 experiments. Cellular fluorescence intensity was expressed as mean integrated density as a function of individual cell size. Percentage colocalization was calculated using the JACoP plugin in single Z-stack sections of deconvolved images. Colocalization is shown in native images and/or as white pixels using the “colocalization finder” plugin in ImageJ, using same threshold for all the images (Schneider et al. 2012).

### Reverse transcription (RT)

Total RNA was extracted using the Tripure™ Isolation Reagent (Roche Applied Science, Mannheim, Germany), according to the manufacturer’s instructions. The yield of total RNA was determined by measuring the absorbance (260/280 nm) using a NanoDropND-1000 spectrophotometer (Nano-Drop Technologies, USA). RT was completed with the high-capacity cDNA Reverse Transcription Kit (Applied Biosystems, Foster City, CA, USA), following manufacturer instructions. Reactions were performed for 10 min at 25°C, 2 h at 37°C and terminated by heating for 5 sec at 85°C. The reaction mixture was maintained at −20°C until further use.

### Quantitative real-time PCR

Quantitative real-time PCR of the different mRNAs was performed in triplicate using gene-specific primers and SYBR® Green. Oligonucleotide primers were designed using Primer Express 2.0 software (Applied Biosystems, Foster City, CA, USA). The primer sequences and corresponding GenBank accession numbers are given in table 1. As an internal control for normalization, PCR reactions were performed concurrently with the amplification of a reference gene, 18S ribosomal RNA (rRNA) that proved to be stable in all the conditions studied.

**Table 1.**
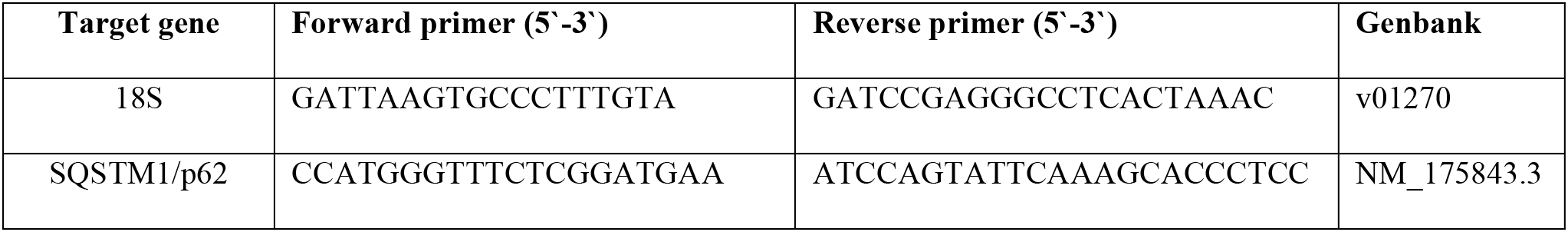
Real time PCR primers the GenBank accession numbers.

Real time-PCR was performed on an Step-one plus (Applied Biosystems, Foster City, CA, USA) real-time thermal cycler using the SYBR® Green PCR Master Mix kit (Applied Biosystems, Foster City, CA, USA) with the following thermal cycler settings: one cycle of 10 min at 95°C, 40 cycles of 15 s at 95°C and 1 min at 60°C. Cycle thresholds for both genes were selected immediately above the baseline and within the linear range on log scaling. Each reaction (20 μl) consisted of a 2 μl cDNA aliquot, 300 nM of each primer and 10 μl of SYBR® Green PCR Master Mix containing AmpliTaq gold DNA polymerase. Increases in the amount of SYBR® Green reporter dye fluorescence during the amplification process were analyzed with Sequence Detector software (SDS version 1.6 Applied Biosystems, Foster City, CA, USA). Relative change in expression of the target genes was determined by the following equation:

Fold change = 2−ΔCt, ΔCt = (Ct target–Ct 18S ribosomal RNA (rRNA)) (Livak and Schmittgen, 2001).

The Ct value is the cycle number at which the fluorescence signal crosses the designated threshold.

### Morphological studies

For ultrastructural studies, HGs were treated as previously described (Vega-Naredo et al. 2009). HGs were lightly fixed by immersion in a solution containing 1.5% glutaraldehyde and 2.5% paraformaldehyde in 0.1 M phosphate buffer (pH 7.4). Fixation was continued overnight at 4°C using fresh fixing solutions. Tissues were then postfixed in 1% Osmium (OsO4) for 2 hours. After dehydration in a graded acetone series, the tissue fragments were embedded in the epoxy resin TAAB 812, and 1 μm semithin sections were stained with toluidine blue. Ultrathin sections were collected on copper grids, stained with uranyl acetate-lead citrate, and examined using a Zeiss EM-109 transmission electron microscope (Zeiss, Oberkochen Germany) operating at 80 kV.

### Mitochondrial functionality

As an indicator of the mitochondrial population, citrate synthase (CS) activity was determined spectrophotometrically at 412 nm and 30°C, as previously described (Garcia-Macia et al. 2011).

The intracellular ATP content in the HGs was measured with an Adenosine-5’-triphosphate bioluminescent assay kit (FL-AA, Sigma Aldrich Inc.). Bioluminescent luciferase-luciferin reactions provide the basis of simple, rapid, and highly sensitive assays for ATP (Taylor et al. 1998). Samples (100 μL) of tissue homogenates diluted 1:100 were mixed with 100 μL of ATP assay mix dilution buffer FL-AAB (pH 7.8). Light production was then immediately measured by luminescence using a Luminometer Turner Designs TD-20/20 (Turner BioSystems Inc., Sunnyvale, CA, USA) (Yen et al. 2007).

### Triglyceride levels

The triglyceride (TG) levels were determined using a triglyceride quantification kit following the manufacturer’s indications (Abcam, Cambridge, UK). In this assay, triglycerides were converted to free fatty acids and glycerol. The glycerol was oxidized and combined with a probe to generate the luminometric signal. HG gland samples were lysed using NP-40 solution, then incubated with the lipase for 20 minutes. A final incubation of 1 hour with the Triglyceride reaction to ensure the oxidation of the glycerol and the colorimetric reaction. Measurement was performed at OD 570 nm in a Varioskan Flash (Thermo Fisher, Waltham, MA, USA).

## RESULTS

### Melatonin receptors expression: cellular response to oxidative stress

The role of melatonin in estrous cycle physiology can be assessed by the expression of melatonin receptors, which have been described in HG (Tomas-Zapico et al. 2005). MT1 is a G protein-coupled receptor at the plasma membrane, and it is activated by the pineal-synthesized melatonin (Tomas-Zapico et al. 2005). RORα is a nuclear melatonin receptor that plays an important role in cell protection against oxidative stress (Caballero et al. 2008).Western-blot analyses for MT_1_ and RORα proteins showed higher expression levels of both in diestrus (Fig. 1a, p<0.05).

**Figure 1.**
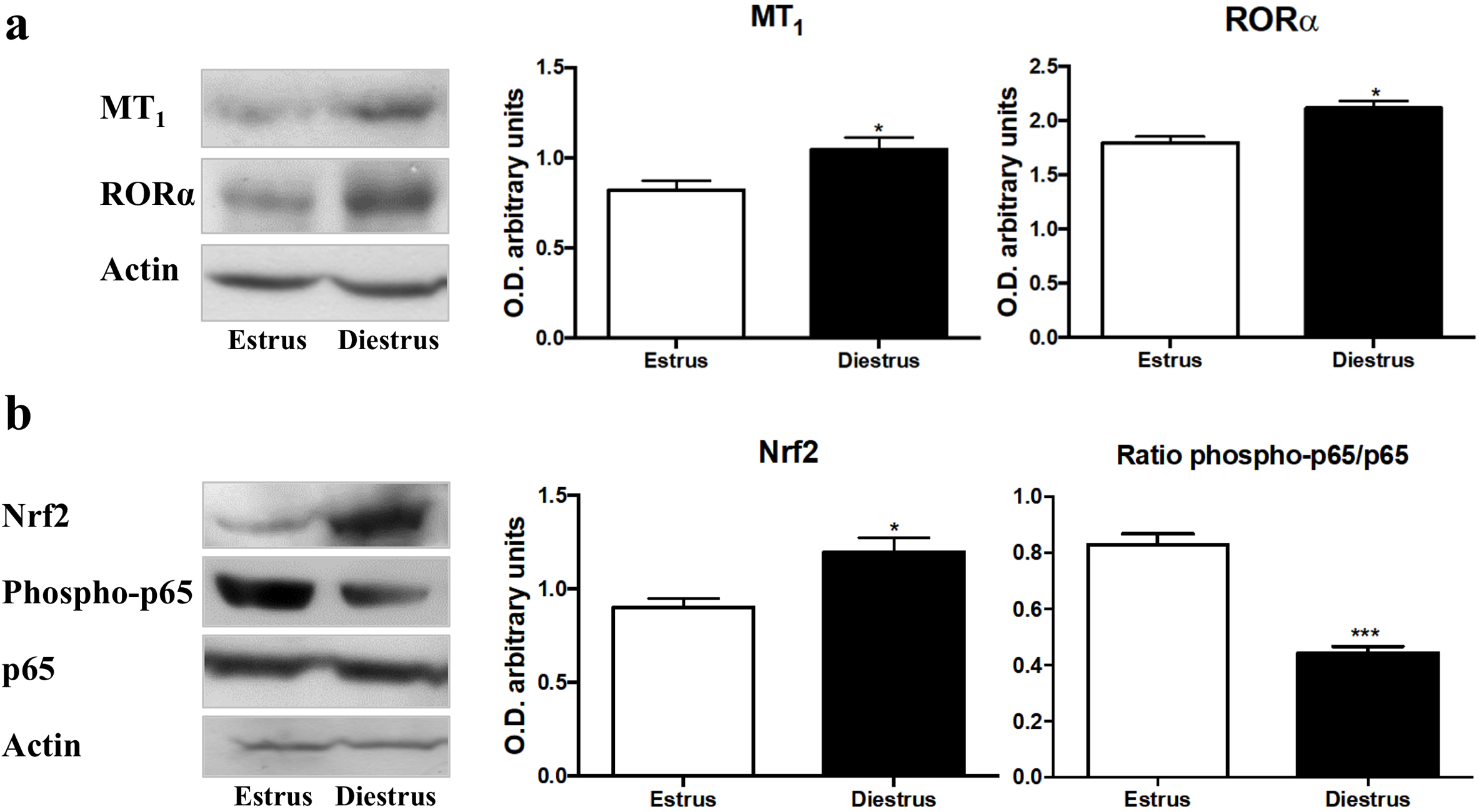
Melatonin role in the cellular response to oxidative stress. **(a)** MT-1, RORα and **(b)** Nrf2, phospo-p65 and p65 protein band intensities in the different fractions of the Harderian glands from the estrus and diestrus phases. The bar graphs of MT-1 and RORα **(a)** and Nrf2 (b) quantifies the optical densities of the western blot bands normalized to ß-actin and the phospho-p65/ p65 ratio **(b)** as NF-κB activation marker.

Nrf2 is an essential transcription factor in the cellular response to oxidative stress (Zhang 2006). Under basal conditions, nuclear levels of Nrf2 remain relatively low because it is sequestered in the cytosol (Negi et al. 2011; Zhang 2006). High oxidative stress levels activate the canonical Nrf2 pathway (Lau et al. 2010). Our study reveals a strong Nrf2 activation in the diestrus phase, compared to the one in estrus phase (Fig. 1b, p<0.05).

Nuclear factor-kappa B (NF-κB) is a transcription factor that also plays a key role in cell responses to oxidative stress (Michiels et al. 2002). We examined its activation state using immunoblot analysis of key protein in the NF-κB pathway: p65 and phospho-p65 (Ser536). Activation occurs via phosphorylation of p65 at Ser536, and only the phosphorylated protein has nuclear localization and transcriptional activity (Vega-Naredo et al. 2009). Total amount of p65 did not show significant differences between phases of the estrous cycle of HG (Fig. 1b). However, phosphorylated p65 expression was significantly lower in diestrus than estrus phase (Fig. 1b). Thus, to characterize the NF-κB activation, we evaluated the ratio of phospho-p65 respect to total p65 protein and confirmed a lower level of phospho-p65/p65 in diestrus compared to estrus (Fig. 1b, p<0.001). Our data suggest a lower activation level of the NF-κB pathway in the diestrus phase.

### Functional mitochondrial status

Mitochondria play an essential role in cell survival (Caballero et al. 2013), but their activity generates free radicals that alter the oxidative balance (Leon et al. 2004). Mitochondria also play a key role in lipid metabolism (Schatz 1995). To characterize mitochondrial functional status, we studied citrate synthase activity and ATP generation. Citrate synthase is a mitochondrial matrix enzyme, essential in Krebs cycle that is an indicator of healthy mitochondrial population (Garcia-Macia et al. 2011). Based on this mitochondrial marker, the number of mitochondria is higher in diestrus than in estrus (Fig. 2a, p<0.05). Accordingly, ATP levels were also higher in diestrus, indicating better mitochondrial efficiency (Fig. 2b, p<0.001).

**Figure 2.**
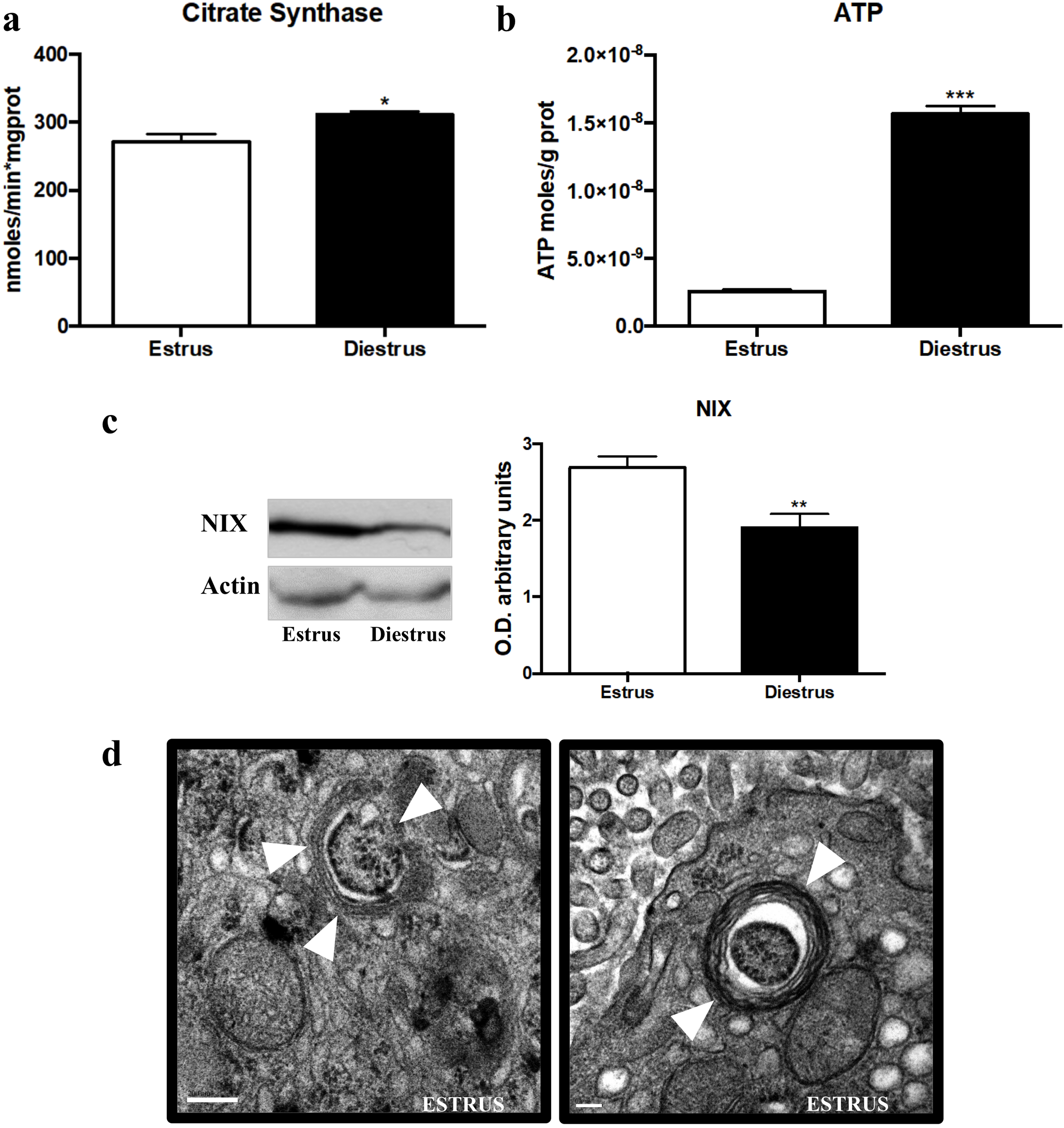
Mitochondrial status and clearance. Mitocondrial **(a)** Comparison of Citrate Synthase (CS) activity, which results are expressed in nmoles/min*mg protein. **(b)** ATP intracellular content, which results are expressed as ATP moles/g protein. **(c)** NIX protein band intensity and NIX bar graph that quantifies the optical densities of the western blot bands normalized to ß-actin. Data are expressed as the means ± SEM and are calculated from at least three separate experiments. *: (p< 0.05), **: (p<0.01), ***: (p<0.001). **(d)** Electron micrographs of female cells in estrus phase showing mitochondria within an autophagosome (arrowhead) as representative images. Similar results were obtained from three separate experiments. Scale bar, 0.2 μm.

Mitophagy, the selective autophagy pathway to degrade defective mitochondria, was also assessed. The mitochondrial protein NIX has been described as an autophagy receptor, mediating the clearance of damaged mitochondria (Novak et al. 2010). NIX expression was lower in diestrus than estrus (Fig. 2c, p<0.01). Using electron microscopy (EM) we corroborated the elevated presence of mitophagosomes during estrus (Fig. 2d). Mitochondria are healthier in diestrus, hence, their selective degradation through autophagy is less prominent during this phase.

### Lipophagy

The LC3-interacting protein SQSTM1/p62 is a key autophagic protein for cell homeostasis mediating the selective specific degradation of protein aggregates and cytoplasmic bodies (Lin et al. 2013). Recent studies have described SQSTM1/p62 as a key mediator in lipolysis (Lee et al. 2010) and also, in lipophagy (Tatsumi et al. 2018; Wang et al. 2017). SQSTM1/p62 protein (Fig. 3a, p<0.01) and RNA (Fig. 3b, p<0.01) levels were significantly lower during the diestrus.

**Figure 3.**
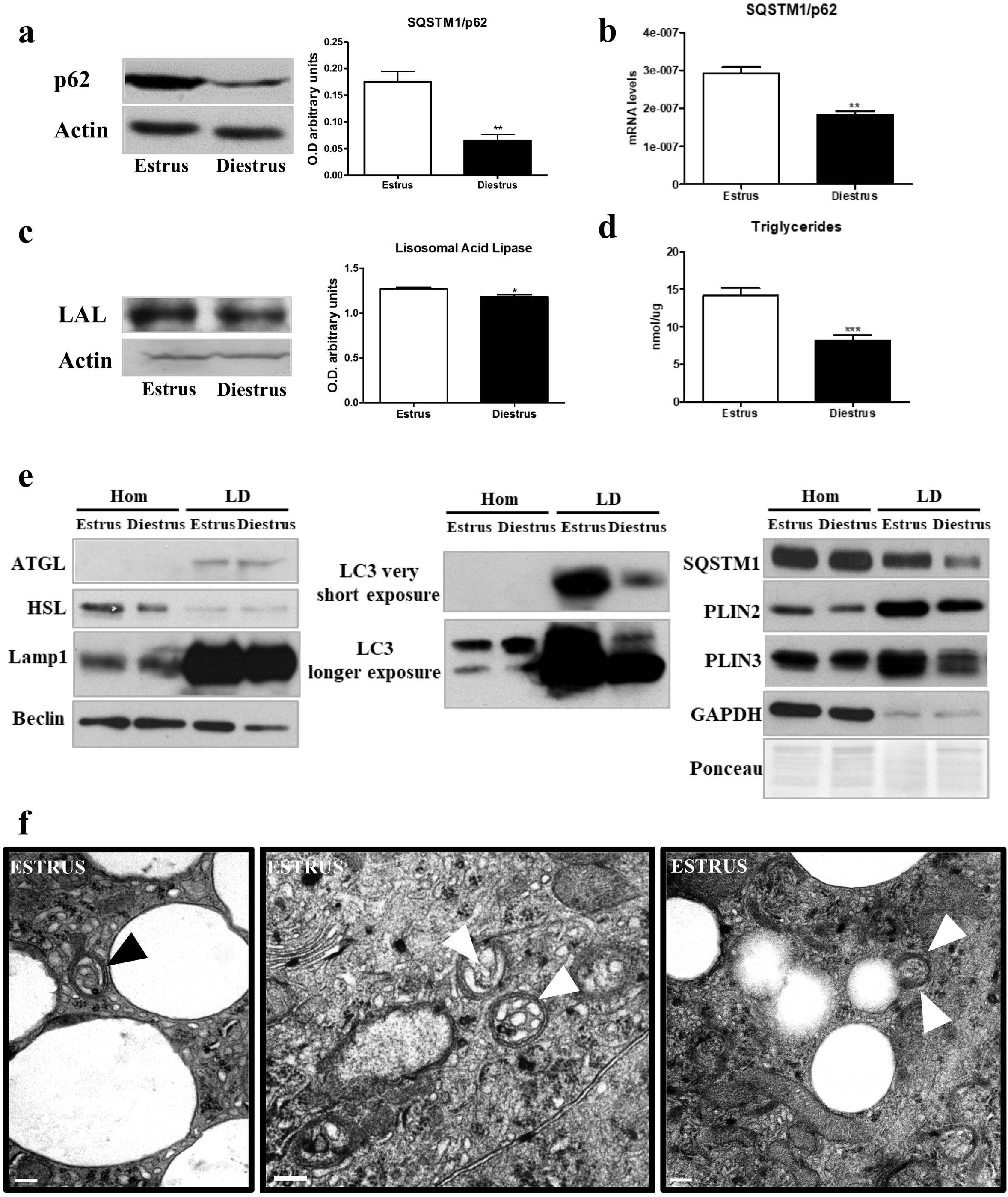
Lipolisis-related autophagy activity. **(a)** SQSTM1/p62 and **(c)** LAL protein band intensities and its respective bar graphs that quantifies the optical densities of the western blot (WB) bands normalized to ß-actin. **(b)** SQSTM1/p62 mRNA expression quantification, normalized to 18s. **(d)** Triglyceride levels in estrus and diestrus, results are expressed in nmoles/μg protein. **(e)** WB for indicated proteins in total homogenates (Hom) and lipid droplets (LD). Data are expressed as the means ± SEM and are calculated from at least three separate experiments, with each experiment performed in triplicate. *: (p< 0.05) **: (p< 0.01) **(f)** Electron micrographs of female cells in estrus phase showing lipid droplets within autophagosomes (arrowheads). Scale bar, 200 nm.

Lipolytic processes were studied by assaying the expression of lysosomal acid lipase (LAL) and the triglyceride (TG) levels. LAL is involved in the degradation of cholesteryl esters and triglycerides (Anderson et al. 1999). Our data showed LAL lower expression in the diestrus phase (Fig. 3c, p<0.05). Concomitantly, TGs levels were also lower in the diestrus phase (Fig. 3d, p<0.001). LAL degrades the lipids provided by autophagy machinery (lipophagy) in the lysosome (Skop et al. 2012). To characterize the level of lipophagy, we quantified the presence of the autophagy machinery components in the LD in both phases. First, we isolated LDs from HGs at estrus and diestrus, the expression of cytosolic lipases (ATGL and HSL) was assessed to evaluate classical lipolysis, no changes were found in the LD isolations. Then, we determined the presence and expression levels of autophagic proteins as Lamp1, Beclin, LC3-II, SQSTM1/p62, Plin2, Plin3 and, also, GAPDH to exclude cytosolic contamination (Martinez-Lopez et al. 2016), by western blot. The expression of all the autophagic proteins was lower in LDs from the diestrus phase while the total homogenate (Hom) was unchanged (Fig. 3e). This result indicates lower LD degradation in diestrus than estrus phase. Furthermore, our results from EM analysis of HGs during the estrous cycle revealed the presence of lipid droplets inside the autophagosomes showing the high levels of lipid degradation (Fig 3f). In order to further quantify the abundance of LDs we analyze the presence of the perilipin (Plin) family proteins that coats LDs (Itabe et al. 2017). As expected, we observed lower expression of Plin2 and Plin3 in diestrus, indicating fewer LDs in this phase (Fig. 3e).

We then use immunohistochemistry to confirm the previous results and determine whether LC3 (Fig. 4a) and Lamp1 (Fig. 4b), autophagosome and lysosomal markers respectively, co-localize with BODIPY, a dye that stains neutral lipids in the LD. We found less colocalization of both markers with the LDs in diestrus phase (Fig 4a&b, p<0.001), supporting the idea of higher lipophagy levels during estrus.

**Figure 4.**
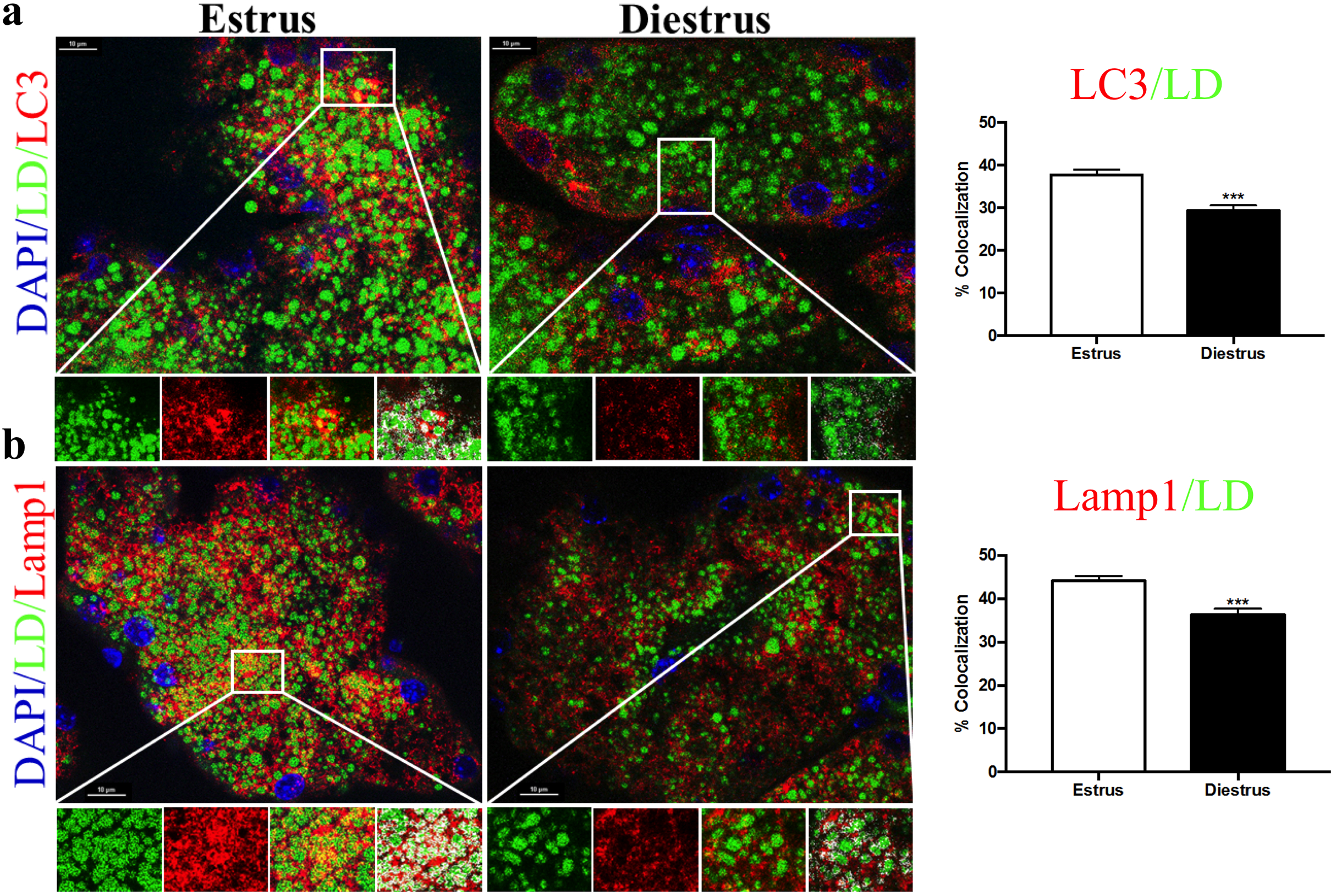
Differential lipophagy activation during the estrous cycle. Immunofluorescence for LC3 and BODIPY 493/503 **(a)** and Lamp1 and BODIPY 493/503. Colocalization determined by the “colocalization finder” plugin in ImageJ is shown in white in the fourth enlarged panel. Enlarged image scale bar: 10μm. **(b)** and its respective bar graphs that quantifies the percentage of colocalization.

## DISCUSION

How sexual hormones modulate selective autophagy is poorly understood, this lack in knowledge aggravates the sexual bias we could easily find in biological research. Harderian gland (HG) has an intrinsic plasticity to quickly respond to several internal and external stimuli via both morphological and biochemical changes (Rodriguez-Colunga et al. 1993). We have previously shown that female HG variations are dependent on the fluctuations in sex hormones during the estrous cycle, which makes HG the perfect model to understand how these hormonal changes modulate autophagy. These autophagic changes during estrous cycle are conserved among rodents (Koenig et al. 2015).

Melatonin is a hormone secreted by the pineal gland, usually related to protection against oxidative stress due to its antioxidant properties (Vega-Naredo et al. 2012). The rate of melatonin secretion varies synchronously with the vaginal cycle, being higher when the estrogen levels are lower at diestrus (Ozaki et al. 1978). It has been described that melatonin displays an anti-estrogenic effect (Rato et al. 1999) mediated by the membrane-bound receptor MT_1_ (Girgert et al. 2009), although the contribution of nuclear receptors is still unclear (Girgert et al. 2009). Accordingly, melatonin receptors, both MT_1_ (plasma membrane) and RORα (nucleus), are more abundant in diestrus. Thus, the increased melatonin signaling, via its receptors, may be involved in reduction of pro-inflammatory mediators and alleviating the higher oxidative stress levels that exist during the estrus phase (Garcia-Macia et al. 2014). In its role as antioxidant, melatonin has been described as inductor of Nrf2 activation, enhancing its nuclear translocation and subsequent antioxidant response (ARE) binding (Aparicio-Soto et al. 2014; Negi et al. 2011). This translocation process has been observed in our study under diestrus phase, what leads us to assume that melatonin antioxidant effect in the HG is mediated by its receptors and potentiated by Nrf2 activation in this phase along with minimal estrogen levels.

The transcription factor NF-κB is a pro-inflammatory factor activated by oxidative stress and inhibited by melatonin (Negi et al. 2011). Here we have shown that NF-κB is activated during estrus phase along with elevated levels of oxidative stress. Thus, activation of NF-κB may favor the pro-inflammatory profile in HG at the estrus phase. It has been also shown that NF-κB promotes mitophagy by SQSTM1/p62 induction (Zhong et al. 2016). Accordingly, in our results female HG showed activation of SQSTM1/p62 in estrus phase. Moreover, regarding mitochondrial clearance, the NIX protein, which is localized in the mitochondrial outer membrane, has been defined as a mitochondrial receptor for mitophagy (Novak et al. 2010). Our present results showed a higher NIX expression in the female HG at the estrus phase than in the diestrus one. Additionally, we have observed the presence of mitophagosomes by EM (Fig. 2d). All together these data shows the activation of mitophagy in the female’s HG during estrus mediated by NF-κB activation. Moreover, mitochondrial depolarization, tested with Citrate synthase activity, and the subsequent mitochondrial ROS generation have been previously linked to NIX ability to directly activate the autophagy machinery via mTOR inhibition (Ding et al. 2010). Consistent with these markers and data, mitochondrial energetic functions, assayed by ATP concentration, were significantly lower in the HGs during the estrus phase. Therefore, increased levels of the mitophagy marker NIX, formation of mitophagosomes, and impaired mitochondrial functionality suggest an intense mitochondrial removal by selective autophagy in the female HG at the estrus phase promoted by NF-κB activation.

The role of autophagy in lipid metabolism has been recently discovered and lipophagy has been shown as the main mechanism to mobilize lipid stores (Singh et al. 2009). Lipophagy was first discovered in liver and, currently, is described in many other tissues (Singh and Cuervo 2012). SQSTM1/p62 participates in the selective removal of organelles, as mitochondria or LDs through autophagy (Tatsumi et al. 2018; Wang et al. 2017). SQSTM1/p62 and other autophagic machinery directly contributes to the mobilization of lipids from LDs to lysosomes, where they get degraded by lipases (Singh and Cuervo 2012). Our data showed higher lipolytic activity in estrus phase due to the higher expression of lysosomal acid lipase (LAL). Accordingly, autophagy and lysosomal proteins, but not cytosolic lipases, were accumulated in the LDs in this phase, showing higher lipophagy activity. Furthermore, our EM studies revealed the presence of LDs within the autophagosomes. Perilipins showed less expression in diestrus, indicating lower amount of LDs. Thus, our data relate lipolytic and lipophagy activities with the cellular changes usually observed in the HG during the estrous cycle and, until now, not well understood. The higher lipophagy activity in estrus could be the process behind the Type II cells disappearance.

We have shown that the HG in the estrous cycle is characterized by important variations in estrogen levels and oxidative stress (Fig. 5), where melatonin may be its primary moderator, in basis of its clear relationship with both factors: it is a well-known antioxidant and has important antiestrogenic effects. Melatonin interacts with 2 of the estrogen-signaling pathways and it decreases circulating levels of estradiol (Gonzalez-Gonzalez et al. 2018; Sanchez-Barcelo et al. 2005). Melatonin levels are reduced in the estrus phase and entail the activation of NF-κB and the elevated oxidative stress, which induces not only macroautophagy and CMA (Garcia-Macia et al. 2014; Poole et al. 1981; Riley and Behrman 1991) but also increases other selective autophagic processes. All associated to a decrease in mitochondrial activity, leading to degradation via mitophagy. Likewise, activation of NF-κB may also lead to lipolytic processes, as suggested by the increase of the autophagic and lysosomal proteins in the LDs and the degradation of large lipid droplets in Type II cells. These degradation processes require a high lysosomal effort, as shown by the increases in LAL expression. Melatonin, via both membrane and nuclear receptors, reduces pro-inflammatory mediators and enhances the expression of Nrf-2 in diestrus phase. Consequently, autophagy is blocked, and porphyrin release is reduced, and the gland returns to a rest period.

**Figure 5.**
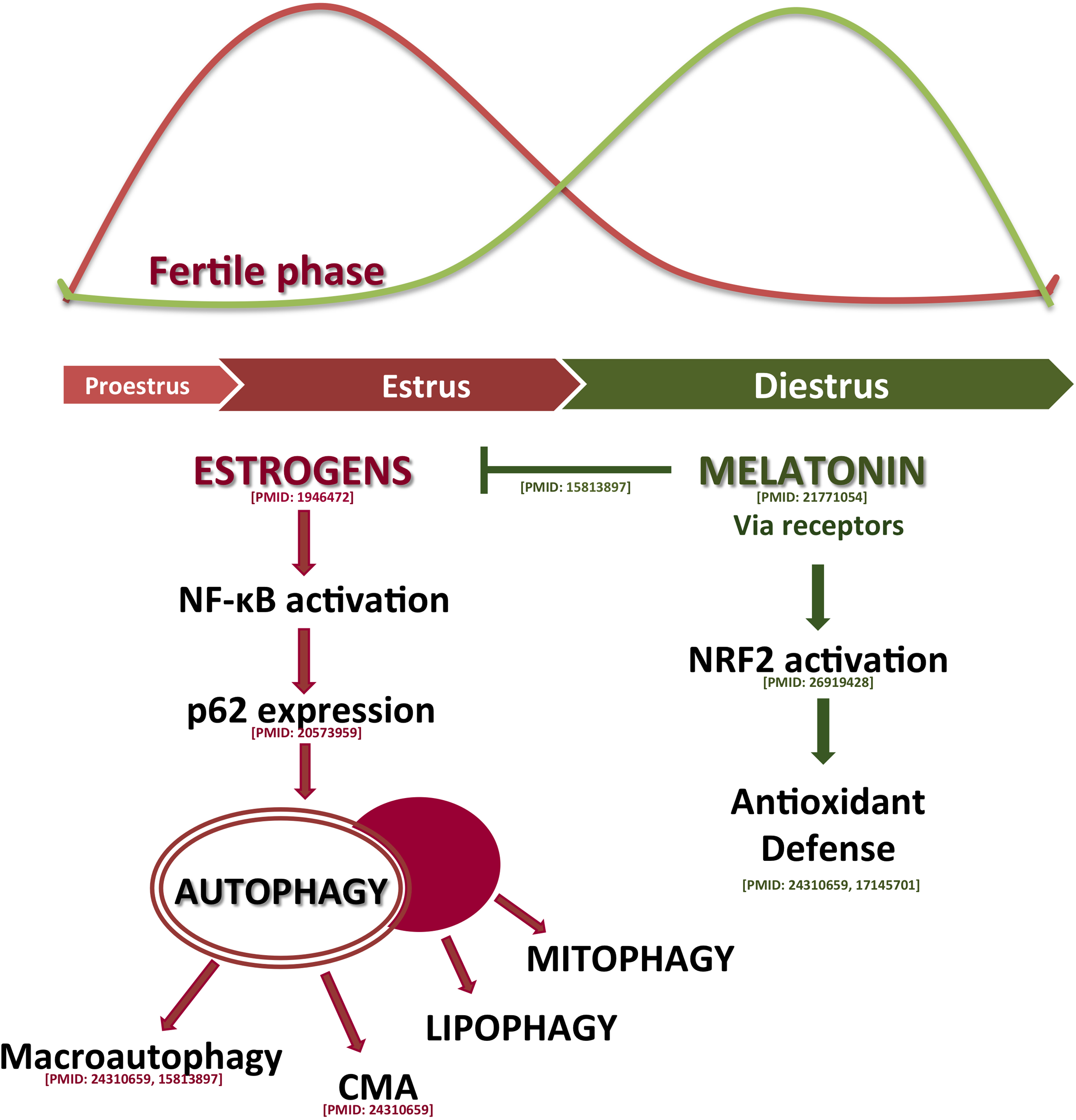
Modulation by melatonin of the estrous cycle mechanisms. The scheme proposes that oxidative stress is one of the key processes involved in the estrous cycle with melatonin as modulator of it.

## Acknowledgments

We are members of the INPROTEOLYS and INEUROPA network. This work was supported by: FISS-18-PI17/02009 and C0245R4032. Y. P has a predoctoral fellowship from the Ministerio de Economia y Competitividad. Financial support from the University of Oviedo is also acknowledged.

## REFERENCES

Anderson RA, Bryson GM, Parks JS (1999) Lysosomal acid lipase mutations that determine phenotype in Wolman and cholesterol ester storage disease Mol Genet Metab Rep. 68:333–345 doi:10.1006/mgme.1999.2904

Aparicio-Soto M, Alarcon-de-la-Lastra C, Cardeno A, Sanchez-Fidalgo S, Sanchez-Hidalgo M (2014) Melatonin modulates microsomal PGE synthase 1 and NF-E2-related factor-2-regulated antioxidant enzyme expression in LPS-induced murine peritoneal macrophages Br J Pharmacol 171:134–144 doi:10.1111/bph.12428

Bradford MM (1976) A rapid and sensitive method for the quantitation of microgram quantities of protein utilizing the principle of protein-dye binding Anal Biochem. 72:248–254

Caballero B, Veenman L, Gavish M (2013) Role of mitochondrial translocator protein (18 kDa) on mitochondrial-related cell death processes Recent Pat Endocr Metab Immune Drug Discov 7:86–101

Caballero B et al. (2008) Favorable effects of a prolonged treatment with melatonin on the level of oxidative damage and neurodegeneration in senescence-accelerated mice J Pineal Res 45:302–311 doi:10.1111/j.1600-079X.2008.00591.xJPI591 [pii]

Coto-Montes A, Boga JA, Rosales-Corral S, Fuentes-Broto L, Tan DX, Reiter RJ (2012) Role of melatonin in the regulation of autophagy and mitophagy: a review Mol Cell Endocrinol 361:12–23 doi:10.1016/j.mce.2012.04.009

Coto-Montes A et al. (2009) Sexual autophagic differences in the androgen-dependent flank organ of Syrian hamsters J Androl 30:113–121 doi:10.2164/jandrol.108.005355

Ding WX et al. (2010) Nix is critical to two distinct phases of mitophagy, reactive oxygen species-mediated autophagy induction and Parkin-ubiquitin-p62-mediated mitochondrial priming J Biol Chem 285:27879–27890 doi:10.1074/jbc.M110.119537

Du L et al. (2009) Starving neurons show sex difference in autophagy J Biol Chem 284:2383–2396 doi:10.1074/jbc.M804396200

Duta-Mare M et al. (2018) Lysosomal acid lipase regulates fatty acid channeling in brown adipose tissue to maintain thermogenesis Biochim Biophys Acta Mol Cell Biol Lipids 1863:467–478 doi:10.1016/j.bbalip.2018.01.011

Garcia-Macia M et al. (2014) Autophagic and proteolytic processes in the Harderian gland are modulated during the estrous cycle Histochem Cell Biol 141:519–529 doi:10.1007/s00418-013-1170-1

Garcia-Macia M et al. (2011) Melatonin induces neural SOD2 expression independent of the NF-kappaB pathway and improves the mitochondrial population and function in old mice J Pineal Res 50:54–63 doi:10.1111/j.1600-079X.2010.00809.x

Girgert R, Hanf V, Emons G, Grundker C (2009) Membrane-bound melatonin receptor MT1 down-regulates estrogen responsive genes in breast cancer cells J Pineal Res 47:23–31 doi:10.1111/j.1600-079X.2009.00684.x

Gonzalez-Gonzalez A, Mediavilla MD, Sanchez-Barcelo EJ (2018) Melatonin: A Molecule for Reducing Breast Cancer Risk Molecules 23 doi:10.3390/molecules23020336

Hoffman RA, Johnson LB, Reiter RJ (1985) Harderian glands of golden hamsters: temporal and sexual differences in immunoreactive melatonin J Pineal Res 2:161–168

Itabe H, Yamaguchi T, Nimura S, Sasabe N (2017) Perilipins: a diversity of intracellular lipid droplet proteins Lipids Health Dis 16:83 doi:10.1186/s12944-017-0473-y

Koenig U et al. (2015) Autophagy facilitates secretion and protects against degeneration of the Harderian gland Autophagy 11:298–313 doi:10.4161/15548627.2014.978221

Lau A et al. (2010) A noncanonical mechanism of Nrf2 activation by autophagy deficiency: direct interaction between Keap1 and p62 Mol Cell Biol 30:3275–3285 doi:10.1128/MCB.00248-10MCB.00248-10 [pii]

Lee SJ et al. (2010) A functional role for the p62-ERK1 axis in the control of energy homeostasis and adipogenesis EMBO Rep 11:226–232 doi:10.1038/embor.2010.7

Leon J, Acuna-Castroviejo D, Sainz RM, Mayo JC, Tan DX, Reiter RJ (2004) Melatonin and mitochondrial function Life Sci 75:765–790 doi:10.1016/j.lfs.2004.03.003S0024320504003236 [pii]

Lin X, Li S, Zhao Y, Ma X, Zhang K, He X, Wang Z (2013) Interaction Domains of p62: A Bridge Between p62 and Selective Autophagy DNA Cell Biol doi:10.1089/dna.2012.1915

Lisk RD (1985) The estrous cycle. In: The hamster: reproduction and behavior. New York: Plenum Press, pp 23–51

Martin M, Macias M, Escames G, Reiter RJ, Agapito MT, Ortiz GG, Acuna-Castroviejo D (2000) Melatonin-induced increased activity of the respiratory chain complexes I and IV can prevent mitochondrial damage induced by ruthenium red in vivo J Pineal Res 28:242–248

Martinez-Lopez N et al. (2016) Autophagy in the CNS and Periphery Coordinate Lipophagy and Lipolysis in the Brown Adipose Tissue and Liver Cell Metab 23:113–127 doi:10.1016/j.cmet.2015.10.008

Menendez-Pelaez A, Lopez-Gonzalez MA, Guerrero JM (1993) Melatonin binding sites in the Harderian gland of Syrian hamsters: sexual differences and effect of castration J Pineal Res 14:34–38

Michiels C, Minet E, Mottet D, Raes M (2002) Regulation of gene expression by oxygen: NF-kappaB and HIF-1, two extremes Free Radic Biol Med 33:1231–1242

Negi G, Kumar A, Sharma SS (2011) Melatonin modulates neuroinflammation and oxidative stress in experimental diabetic neuropathy: effects on NF-kappaB and Nrf2 cascades J Pineal Res 50:124–131 doi:10.1111/j.1600-079X.2010.00821.x

Novak I et al. (2010) Nix is a selective autophagy receptor for mitochondrial clearance EMBO Rep 11:45–51 doi:10.1038/embor.2009.256

Orsini MW (1961) The external vaginal phenomena characterizing the stages of the estrous cycle, pregnancy, pseudopregnancy, lactation, and the anestrous hamster, *Mesocricetus auratus* waterhouse Lab Anim Care 11:193–206

Ozaki Y, Wurtman RJ, Alonso R, Lynch HJ (1978) Melatonin secretion decreases during the proestrous stage of the rat estrous cycle Proc Natl Acad Sci U S A 75:531–534

Payne AP, McGadey J, Moore MH, Thompson GG (1979) Changes in Harderian gland activity in the female golden hamster during the oestrous cycle, pregnancy and lactation Biochem J. 178:597–604

Pearson GL et al. (2014) Lysosomal acid lipase and lipophagy are constitutive negative regulators of glucose-stimulated insulin secretion from pancreatic beta cells Diabetologia 57:129–139 doi:10.1007/s00125-013-3083-x

Poole MC, Mahesh VB, Costoff A (1981) Morphometric analysis of the autophagic and crinophagic lysosomal systems in mammotropes throughout the estrous cycle of the rat Cell Tissue Res 220:131–137

Rato AG, Pedrero JG, Martinez MA, del Rio B, Lazo PS, Ramos S (1999) Melatonin blocks the activation of estrogen receptor for DNA binding FASEB J 13:857–868

Reiter RJ (1980) The pineal and its hormones in the control of reproduction in mammals Endocr Rev 1:109–131

Riley JC, Behrman HR (1991) Oxygen radicals and reactive oxygen species in reproduction Proc Soc Exp Biol Med 198:781–791

Rodriguez-Colunga MJ, Rodriguez C, Antolin I, Uria H, Tolivia D, Vaughan MK, Menendez-Pelaez A (1993) Development and androgen regulation of the secretory cell types of the Syrian hamster (Mesocricetus auratus) Harderian gland Cell Tissue Res 274:189–197

Sakai T (1981) The mammalian Harderian gland: morphology, biochemistry, function and phylogeny Arch Histol Jpn 44:299–333

Sanchez-Barcelo EJ, Cos S, Mediavilla D, Martinez-Campa C, Gonzalez A, Alonso-Gonzalez C (2005) Melatonin-estrogen interactions in breast cancer J Pineal Res 38:217–222 doi:10.1111/j.1600-079X.2004.00207.x

Schatz G (1995) Mitochondria: beyond oxidative phosphorylation Biochim Biophys Acta 1271:123–126

Schneider CA, Rasband WS, Eliceiri KW (2012) NIH Image to ImageJ: 25 years of image analysis Nat Methods 9:671–675

Singh R, Cuervo AM (2012) Lipophagy: connecting autophagy and lipid metabolism Int J Cell Biol 2012:282041 doi:10.1155/2012/282041

Singh R et al. (2009) Autophagy regulates lipid metabolism Nature 458:1131–1135 doi:10.1038/nature07976

Skop V et al. (2012) Autophagy-lysosomal pathway is involved in lipid degradation in rat liver Physiol Res 61:287–297 doi:932285 [pii]

Tamarindo GH et al. (2019) Melatonin and Docosahexaenoic Acid Decrease Proliferation of PNT1A Prostate Benign Cells via Modulation of Mitochondrial Bioenergetics and ROS Production Oxid Med Cell Longev 2019:5080798 doi:10.1155/2019/5080798

Tatsumi T et al. (2018) Forced lipophagy reveals that lipid droplets are required for early embryonic development in mouse Development 145 doi:10.1242/dev.161893

Taylor AL, Kudlow BA, Marrs KL, Gruenert DC, Guggino WB, Schwiebert EM (1998) Bioluminescence detection of ATP release mechanisms in epithelia Am J Physiol 275:C1391–1406

Tomas-Zapico C et al. (2005) Coexpression of MT1 and RORalpha1 melatonin receptors in the Syrian hamster Harderian gland J Pineal Res 39:21–26

Vega-Naredo I et al. (2012) Melatonin modulates autophagy through a redox-mediated action in female Syrian hamster Harderian gland controlling cell types and gland activity J Pineal Res 52:80–92 doi:10.1111/j.1600-079X.2011.00922.x

Vega-Naredo I et al. (2009) Sexual dimorphism of autophagy in Syrian hamster Harderian gland culminates in a holocrine secretion in female glands Autophagy 5:1004–1017

Wang L, Zhou J, Yan S, Lei G, Lee CH, Yin XM (2017) Ethanol-triggered Lipophagy Requires SQSTM1 in AML12 Hepatic Cells Sci Rep 7:12307 doi:10.1038/s41598-017-12485-2

Yen CC, Lu FJ, Huang CF, Chen WK, Liu SH, Lin-Shiau SY (2007) The diabetogenic effects of the combination of humic acid and arsenic: in vitro and in vivo studies Toxicol Lett 172:91–105 doi:10.1016/j.toxlet.2007.05.008

Zhang DD (2006) Mechanistic studies of the Nrf2-Keap1 signaling pathway Drug Metab Rev 38:769–789 doi:10.1080/03602530600971974

Zhong Z et al. (2016) NF-kappaB Restricts Inflammasome Activation via Elimination of Damaged Mitochondria Cell 164:896–910 doi:10.1016/j.cell.2015.12.057

Zielniok K, Motyl T, Gajewska M (2014) Functional interactions between 17 beta-estradiol and progesterone regulate autophagy during acini formation by bovine mammary epithelial cells in 3D cultures Biomed Res Int 2014:382653 doi:10.1155/2014/382653

Zielniok K, Sobolewska A, Gajewska M (2017) Mechanisms of autophagy induction by sex steroids in bovine mammary epithelial cells J Mol Endocrinol 59:29–48 doi:10.1530/JME-16-0247

